# AGPI: An AI-Powered Genomic Pathogen Intelligence Platform for Integrated Classification, Visualization, and Therapeutic Targeting

**DOI:** 10.64898/2026.07.07.737037

**Authors:** Aadya Goel, Pallavi Mishra

## Abstract

Rapid and accurate pathogen detection remains a major challenge in modern bioinformatics, as existing tools are often fragmented and require multiple specialized workflows. We present AGPI (AI-powered Genomic Pathogen Intelligence), an integrated platform that combines genomic sequence classification, biological enrichment, three-dimensional structural visualization, and AI-guided therapeutic prioritization within a single interpretable pipeline. AGPI employs a hybrid convolutional–Bidirectional Gated Recurrent Unit (BiGRU) architecture trained on DNA sequences from 40 pathogen classes spanning viruses, bacteria, fungi, and protozoan pathogens. The model achieved 99.61% validation accuracy and 94.90% accuracy on an independent held-out evaluation of 600 pathogen sequences following iterative refinement. As a proof of concept, AGPI correctly classified a Zika virus genome with 96.14% confidence, retrieved curated biological context from 245 peer-reviewed studies, and identified Ribavirin as a leading therapeutic candidate against the Zika NS5 polymerase through AI-guided molecular docking. Multi-metric ligand similarity analysis further differentiated candidate compounds according to their structural and pharmacological properties. These results demonstrate that integrated AI-driven genomic pipelines can accelerate pathogen characterization and therapeutic hypothesis generation while providing an accessible and interpretable framework for infectious disease surveillance and computational drug repurposing.

## Introduction

The emergence of novel infectious pathogens and the rapid evolution of existing ones pose sustained challenges for global public health. Genomic surveillance—the systematic monitoring of pathogen genomes to track diversity, mutations, and spread—is now recognized as a cornerstone of pandemic preparedness and clinical response. Yet despite the proliferation of sequencing technologies and public genomic repositories, translating raw sequence data into actionable diagnostic and therapeutic insights remains laborious. Existing bioinformatics workflows are fragmented across specialist tools: genome classifiers, structural visualizers, mutation analyzers, and docking platforms must each be configured, run, and interpreted independently, imposing a significant expertise barrier for clinical researchers and public health practitioners.

Prior work has demonstrated the individual utility of deep learning approaches across these tasks. Recurrent and convolutional architectures (1) have been applied to viral and bacterial genome classification, with tools such as DeepVirFinder (2) and VirNet (3) achieving strong performance on virus identification from metagenomic sequences. Structure-based drug discovery has been advanced by diffusion-based docking models (4) and deep learning scoring functions (5). Molecular fingerprint methods, including Tanimoto similarity on Morgan and MACCS keys remain widely used for ligand candidate selection (6). Transformer-based architectures applied to genomic sequences, such as DNABERT (7), and protein language models such as ESM (8) have further expanded the toolkit for sequence-level and structure-level analysis.

However, these approaches have largely been developed and evaluated as independent computational tools, each addressing only a specific stage of the pathogen analysis pipeline. Consequently, researchers often rely on multiple software packages to perform sequence classification, structural analysis, biological interpretation, and therapeutic screening, increasing analytical complexity and reducing workflow reproducibility. This fragmentation presents a substantial barrier for researchers seeking an integrated view of pathogen biology, particularly in time-sensitive outbreak investigations. To our knowledge, no previously published platform combines multi-class genomic pathogen classification, automated biological enrichment, interactive three-dimensional structural visualization, and AI-guided therapeutic prioritization (9) within a single end-to-end, interpretable workflow.

Here we introduce AGPI (AI-powered Genomic Pathogen Intelligence), a modular platform designed to address this gap. AGPI unifies four core capabilities: (i) AI-driven pathogen classification using a hybrid Conv1D–BiGRU architecture; (ii) automated biological enrichment retrieving symptoms, virulence factors, immune evasion proteins, and curated PDB and PubChem recommendations; (iii) interactive 3D genome visualization via NVIDIA EsmFold and Biopython; and (iv) protein-ligand docking and multi-metric ligand similarity scoring via DiffDock and GNINA. The platform is designed for real-time processing of large-scale genomic datasets and is delivered through a web-based interface accessible to researchers and clinicians without specialist bioinformatics training. We describe the system architecture and methodology, present validation results across 40 pathogen classes, and demonstrate AGPI’s end-to-end capabilities using a Zika virus genomic sequence as a proof-of-concept case study. Figure 1 illustrates the end-to-end AGPI workflow.

**Fig. 1.**
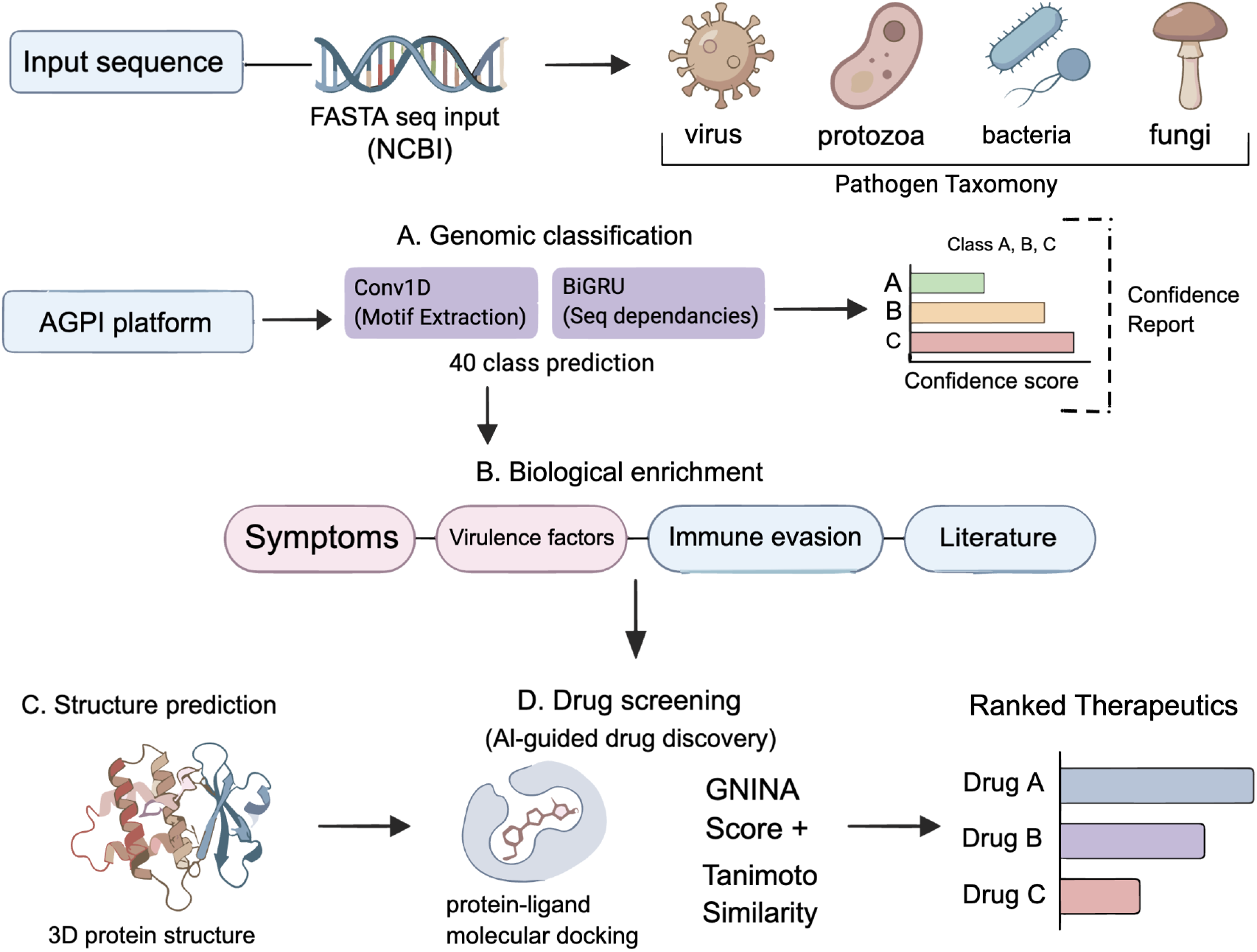
Overview of the AGPI platform pipeline. (A) Genomic classification: input FASTA sequences retrieved from NCBI are classified across 40 pathogen classes spanning viruses, protozoa, bacteria, and fungi using a hybrid Conv1D–BiGRU architecture, with per-class confidence scores reported. (B) Biological enrichment: the predicted pathogen class triggers automated retrieval of associated symptoms, virulence factors, immune evasion mechanisms, and curated literature. (C) Structure prediction: user-input DNA sequences are used to predict 3D protein structures via NVIDIA EsmFold. (D) AI-guided drug screening: predicted structures undergo diffusion-based protein-ligand docking scored by GNINA affinity and Tanimoto-based ligand similarity, yielding a ranked list of therapeutic candidates.

## Results

### Dataset construction and preprocessing

DNA sequences for 40 pathogen classes were retrieved from the NCBI database using BLASTn queries in FASTA format. The dataset spans three biological kingdoms: viruses (including RNA and DNA viruses such as Zika, Dengue, Ebola, SARS-CoV-2, and Hepatitis), bacteria (including *Vibrio cholerae* and other gram-positive and gram-negative species), and fungal pathogens. Pathogens were selected according to three criteria: clinical relevance to global disease burden, diversity of geographic and biological origin, and broad representation across major viral and bacterial families. This selection strategy ensures that the classifier must generalize across both closely related and taxonomically distant organisms. Sequence length varied substantially across classes, ranging from a minimum of 31 nucleotides (Malaria) to a maximum of 7,481 nucleotides (Poliovirus), with per-class mean lengths spanning 100.3 nucleotides (Chickenpox) to 5,000.2 nucleotides (Meningitidis), reflecting the natural diversity of pathogen genome sizes across kingdoms (Table 1). Classes with high intra-class length variance—such as Parvovirus (31–5,243 nt), Poliovirus (877–7,481 nt), and Tuberculosis (36–5,000 nt)—likely contain sequences from multiple genomic regions or strains, which may increase classification difficulty relative to classes with uniform lengths such as Anthrax (1,923 nt) and Zika (3,083 nt).

**Table 1.** Per-class sequence length statistics for the 40 pathogen classes included in the AGPI training corpus. Lengths reported in nucleotides after exact-match deduplication. All sequences were truncated or zero-padded to 1,000 nucleotides for model input. RNA virus sequences were retrieved as DNA-form entries from NCBI GenBank.

| Pathogen | Kingdom | Genome Type | Min | Max | Mean |
| --- | --- | --- | --- | --- | --- |
| Anthrax | Bacteria | dsDNA | 1923 | 1923 | 1923.0 |
| Bocavirus | Virus | ssDNA | 1957 | 1957 | 1957.0 |
| Candidiasis | Fungi | dsDNA | 564 | 1193 | 993.9 |
| Chickenpox | Virus | dsDNA | 41 | 125 | 100.3 |
| Cholera | Bacteria | dsDNA | 1404 | 1424 | 1418.2 |
| Dengue | Virus | ssRNA(+) | 1580 | 1682 | 1669.4 |
| Ebola | Virus | ssRNA(−) | 2406 | 2407 | 2406.2 |
| HFRS | Virus | ssRNA(−) | 1696 | 1697 | 1696.0 |
| HIV | Virus | ssRNA(+)/RT | 34 | 796 | 390.2 |
| Hepatitis | Virus | dsDNA/ssRNA | 1721 | 1727 | 1725.9 |
| Herpes | Virus | dsDNA | 1594 | 1599 | 1597.8 |
| Influenza | Virus | ssRNA(−) | 1494 | 1498 | 1497.5 |
| Legionnaires | Bacteria | dsDNA | 1244 | 1730 | 1642.8 |
| Leishmaniasis | Protozoan | dsDNA | 1182 | 1324 | 1306.7 |
| Leprosy | Bacteria | dsDNA | 230 | 1457 | 566.1 |
| Leptospirosis | Bacteria | dsDNA | 36 | 1485 | 577.3 |
| Lyme | Bacteria | dsDNA | 104 | 1861 | 663.7 |
| Malaria | Protozoan | dsDNA | 31 | 1247 | 358.7 |
| Measles | Virus | ssRNA(−) | 1753 | 1755 | 1755.0 |
| Meningitidis | Bacteria | dsDNA | 5000 | 5001 | 5000.2 |
| Norovirus | Virus | ssRNA(+) | 4468 | 4891 | 4873.1 |
| Papilloma | Virus | dsDNA | 1542 | 2915 | 2511.9 |
| Parvovirus | Virus | ssDNA | 31 | 5243 | 2281.5 |
| Plague | Bacteria | dsDNA | 5000 | 5000 | 5000.0 |
| Poliovirus | Virus | ssRNA(+) | 877 | 7481 | 3176.5 |
| Rabies | Virus | ssRNA(−) | 1571 | 1575 | 1573.4 |
| Rhinovirus | Virus | ssRNA(+) | 44 | 3413 | 2595.4 |
| Rotavirus | Virus | dsRNA | 3240 | 3293 | 3281.0 |
| Salmonellosis | Bacteria | dsDNA | 123 | 5000 | 2225.0 |
| SARS-CoV-2 | Virus | ssRNA(+) | 4254 | 4293 | 4291.3 |
| Smallpox | Virus | dsDNA | 1271 | 1352 | 1326.8 |
| Syphilis | Bacteria | dsDNA | 4973 | 4974 | 4973.4 |
| Torque teno | Virus | ssDNA | 37 | 2659 | 518.6 |
| Tuberculosis | Bacteria | dsDNA | 36 | 5000 | 903.5 |
| UTI | Bacteria | dsDNA | 53 | 4998 | 1743.7 |
| Variola virus | Virus | dsDNA | 4209 | 4209 | 4209.0 |
| Yellow fever | Virus | ssRNA(+) | 3113 | 3226 | 3205.5 |
| Zika | Virus | ssRNA(+) | 3083 | 3083 | 3083.0 |

dsDNA: double-stranded DNA; ssDNA: single-stranded DNA; ssRNA(+/-): positive/negative-sense single-stranded RNA; dsRNA: double-stranded RNA; RT: reverse transcriptase. RNA virus entries retrieved as DNA-form sequences from NCBI GenBank.

Preprocessing followed a standardized pipeline. Sequences were loaded from CSV files and deduplicated by exact sequence match; unique integer class labels were assigned for supervised learning. Nucleotide bases (A, T, C, G) were converted to one-hot encoded binary vectors. To ensure consistent input dimensions, sequences were either truncated or zero-padded to a fixed length of 1,000 nucleotides. The dataset was split into training (80%) and validation (20%) subsets using stratified random sampling to preserve class balance across splits. Crucially, deduplication was performed prior to splitting to prevent identical sequences from appearing in both subsets. The held-out evaluation set of 600 sequences was drawn separately after the train/validation split was finalized, ensuring no overlap with training data. We acknowledge that sequences from the same species or closely related strains may share high nucleotide similarity; future work should apply CD-HIT (10) or similar clustering tools to enforce a minimum sequence identity threshold across splits, which would provide a more conservative estimate of generalization performance. The fixed 1,000-nucleotide input window introduces asymmetric preprocessing effects across classes: pathogens with mean sequence lengths substantially below 1,000 nucleotides—including Chickenpox (mean 100.3 nt), Malaria (mean 358.7 nt), and HIV (mean 390.2 nt)—are heavily zero-padded, while those with sequences exceeding this threshold—including Poliovirus (max 7,481 nt), Meningitidis (max 5,001 nt), and Parvovirus (max 5,243 nt)—undergo truncation that may discard biologically informative coding regions. These preprocessing trade-offs should be considered when interpreting per-class classification performance.

### Hybrid Conv1D–BiGRU classifier achieves near-perfect accuracy on 40 pathogen classes

AGPI’s classification module employs a hybrid architecture combining convolutional and recurrent layers. The model begins with Conv1D layers that extract local sequence features using ReLU activation and max-pooling for dimensionality reduction, with dropout applied for regularization. These local representations are passed to a Bidirectional GRU (BiGRU) layer, which processes the sequence in both forward and backward directions simultaneously to capture long-range upstream and downstream nucleotide dependencies—a critical capability for biological sequences where regulatory elements and coding regions may be spatially separated. The Bi-GRU output feeds into dense layers, culminating in a softmax output over 40 pathogen classes. Training used categorical cross-entropy loss with the Adam optimizer; early stopping on validation accuracy was applied to prevent overfitting.

The model achieved a training accuracy of 99.82% and a validation accuracy of 99.61%, indicating strong generalization with minimal overfitting (Figure 2). These results compare favourably with prior deep learning approaches (11, 12) for genomic sequence classification (2, 3), and the bidirectional context captured by the BiGRU is consistent with established advantages of bidirectional architectures for biological sequence modeling (13).

**Fig. 2.**
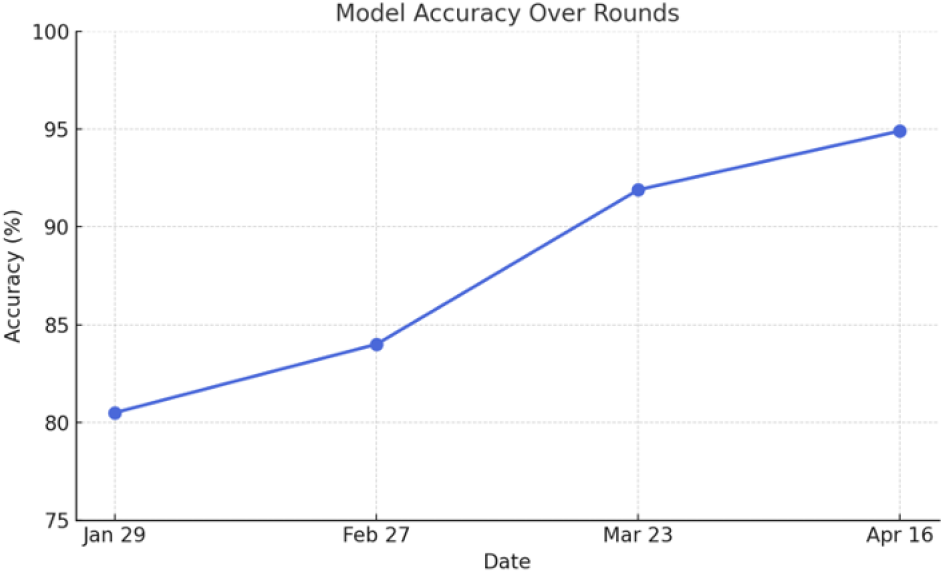
Classification accuracy of the AGPI BiGRU model over four iterative refinement rounds, evaluated on 600 randomly selected pathogen sequences (15 per class across 40 classes). Accuracy improved from 80.50% in Round 1 (January 29) to 94.90% in Round 4 (April 16) following iterative fine-tuning on misclassified samples.

### Iterative held-out evaluation demonstrates robust diagnostic performance

To assess diagnostic performance on unseen sequences, we conducted iterative held-out evaluation on 600 pathogen sequences spanning all 40 classes (15 sequences per class). Each sequence was evaluated independently through the classification pipeline, with predicted class and confidence score logged in a structured evaluation document. Performance was assessed across four iterative rounds, with model fine-tuning targeting misclassified samples between rounds:

- Round 1 (January 29): 80.50%
- Round 2 (February 27): 84.00%
- Round 3 (March 23): 91.89%
- Round 4 (April 16): 94.90%

The steady improvement across rounds confirms that iterative fine-tuning on misclassified samples is an effective strategy for improving diagnostic robustness. The final accuracy of 94.90% across 40 diverse pathogen classes demonstrates that AGPI can serve as a reliable first-pass diagnostic tool for genomic surveillance (Figure 2).

Per-class accuracy across all 40 pathogen classes is shown in Figure 3. All classes exceeded 91% accuracy, with Zika virus achieving the highest per-class accuracy (~ 99%) and Bocavirus the lowest (~ 91%). The narrow performance range across taxonomically diverse pathogens (91–99%) indicates consistent generalization rather than high aggregate accuracy driven by a subset of easier classes—a common failure mode in imbalanced multi-class classifiers. The narrow per-class accuracy range across taxonomically diverse pathogens also carries biological significance. The consistently high performance across both RNA viruses (Zika, Dengue, Ebola) and phylogenetically distinct bacteria and fungi suggests that the hybrid Conv1D–BiGRU architecture successfully captures kingdom-level and family-level nucleotide compositional signatures that are preserved across sequences of the same pathogen class. The comparatively lower accuracy observed for Bocavirus (~91%) may reflect its compact single-stranded DNA genome and high sequence variability among bocaparvovirus strains, which reduces intra-class sequence consistency and increases the likelihood of misclassification at fixed input length. These findings suggest that classifier performance is partly governed by the degree of within-class genomic diversity and genome size relative to the fixed 1,000-nucleotide input window.

**Fig. 3.**
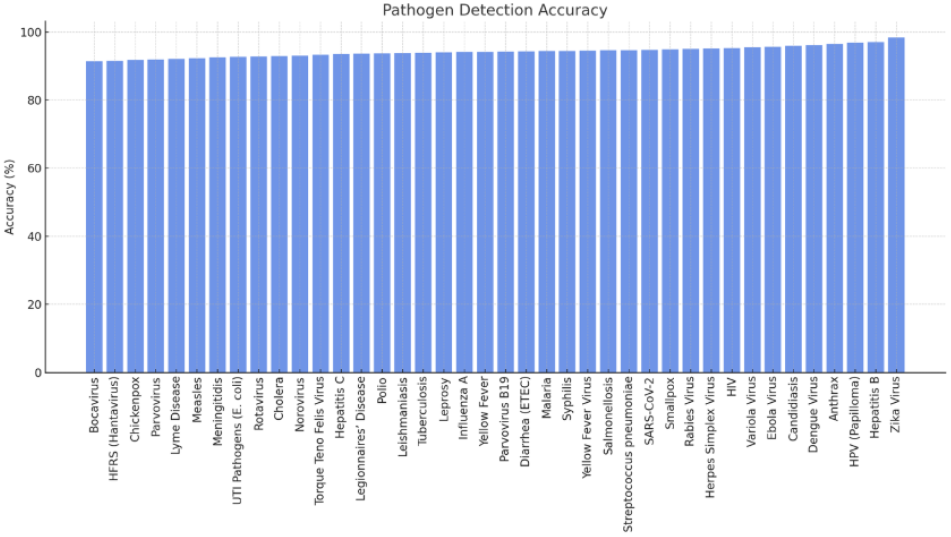
Per-class pathogen detection accuracy of the AGPI classifier across all 40 pathogen classes, sorted by ascending accuracy. All classes exceed 91% accuracy. Zika virus achieves the highest per-class accuracy (~99%); Bocavirus the lowest (~91%). The narrow accuracy range across taxonomically diverse organisms demonstrates consistent model generalization.

### Zika virus case study: end-to-end workflow

To illustrate AGPI’s practical utility, we executed the complete pipeline using a Zika virus genomic sequence as input. The classifier correctly identified the sequence as Zika virus with a confidence score of 96.14%. The biological enrichment module subsequently retrieved contextual information including associated clinical symptoms (fever, rash, conjunctivitis, arthralgia), known virulence factors, functional proteins involved in immune evasion, and a curated list of relevant PDB structures and PubChem compounds drawn from 245 peer-reviewed studies. This enrichment step is designed to immediately orient researchers toward biologically and clinically relevant downstream analyses without requiring manual database queries. The biological enrichment output for Zika virus is clinically coherent: the retrieved symptoms (fever, rash, conjunctivitis, arthralgia) correspond to the established clinical presentation of Zika virus disease, while the identified virulence factors and immune evasion proteins reflect known mechanisms by which Zika evades innate immune responses, including NS5-mediated STAT2 degradation. This correspondence between AGPI’s enrichment output and the peer-reviewed literature provides qualitative evidence that the curated corpus is both accurate and biologically relevant for the tested pathogen class.

### Interactive 3D genome visualization

The AGPI visualization module renders protein structures predicted from user-input DNA sequences using NVIDIA EsmFold (14), a protein language model capable of high-accuracy atomic-level structure prediction. Annotation and interactive rendering are implemented using Biopython (15). For the Zika case study, the full genomic sequence was rendered as an interactive 3D genome browser with annotated, clickable coding regions and functional domains (Figure 4). Researchers can inspect residue-level structural features, track mutation hotspots, and explore protein-ligand binding pockets interactively—capabilities that complement dedicated structural visualization tools such as PyMOL (16) while remaining accessible within a unified web-based pipeline.

**Fig. 4.**
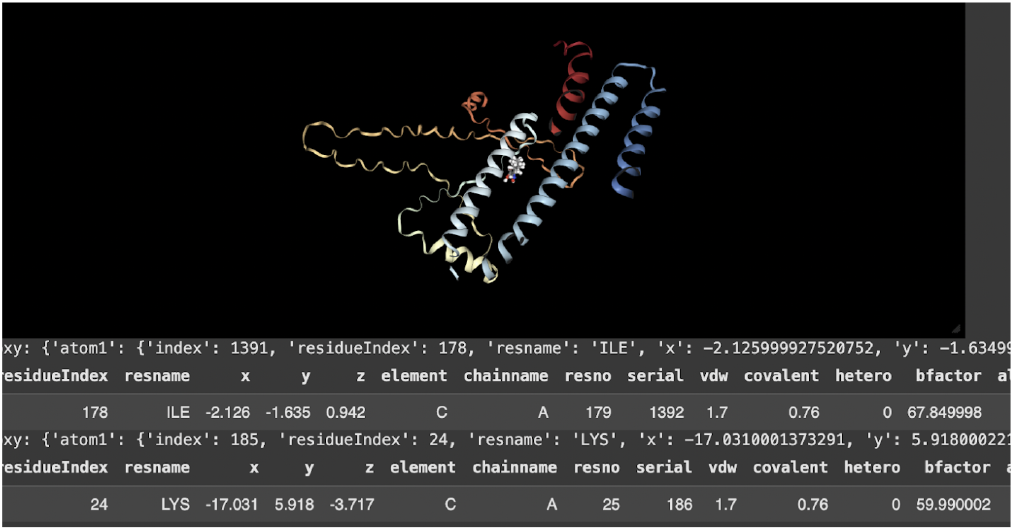
Interactive 3D visualization of the Zika virus genome generated by AGPI using NVIDIA EsmFold for structure prediction and Biopython for annotation. The genome browser displays annotated, clickable coding and functional regions. Residue-level structural data, including binding pocket geometry, are accessible interactively.

### DiffDock protein-ligand docking identifies Ribavirin as a top therapeutic candidate

For therapeutic target identification, AGPI integrates DiffDock (4), a diffusion-based protein-ligand docking model that explicitly models conformational flexibility—a significant advance over traditional rigid-body docking approaches. The docking simulation used the Zika NS5 polymerase (PDB ID: 4QTB) as the target protein and Ribavirin (retrieved by PubChem CID, with SMILES automatically resolved) as the candidate ligand. DiffDock generated 20 binding poses, each scored by both its internal confidence metric and GNINA affinity scores for independent validation.

Ribavirin was ranked as the top therapeutic candidate across poses (Figure 6), with the following top-pose metrics:

- DiffDock Confidence: −1.4
- GNINA Raw Affinity: 30.24
- GNINA Minimized Affinity: −7.31

Correlation analysis across all 20 poses revealed a moderate negative relationship between DiffDock confidence and GNINA scored affinity (*r* = − 0.522), consistent with both metrics capturing energetically favourable binding (Figure 5). The correlation with GNINA minimized affinity was weak (*r* = 0.036), reflecting that post-docking energy minimization introduces conformational adjustments that partially decouple the minimized score from the initial docking confidence. These correlation patterns are informative: they suggest Diff-Dock confidence is a reasonable first-pass ranking criterion, while GNINA minimized affinity provides complementary information and should be considered alongside rather than as a direct substitute. Three-dimensional interaction analysis of the top-ranked pose revealed hydrogen bonds, hydrophobic contacts, and *π*–*π* stacking interactions at the NS5 polymerase active site—a region critical to viral replication. These findings support Ribavirin’s potential for repurposing in Zika treatment and are consistent with prior reports of its broad-spectrum antiviral activity.

**Fig. 5.**
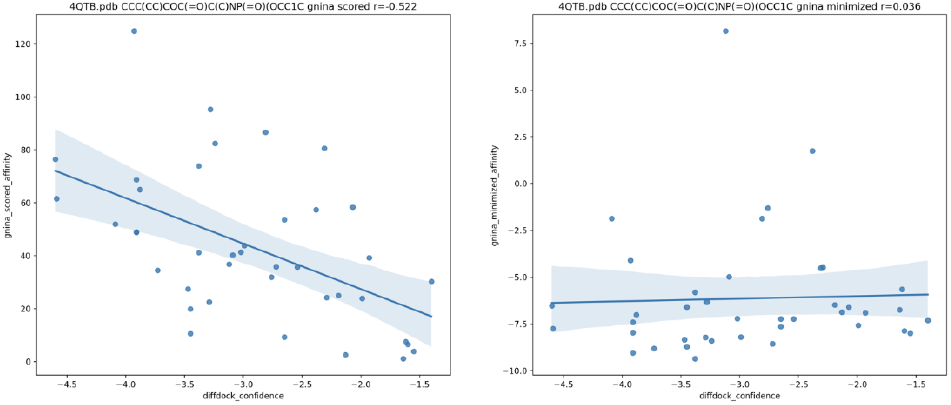
Correlation between DiffDock confidence scores and GNINA affinities across 20 binding poses for Ribavirin docked to the Zika NS5 polymerase (PDB: 4QTB). Left: DiffDock confidence vs. GNINA scored affinity (*r* = −0.522), showing a moderate negative relationship (more negative DiffDock confidence corresponds to higher scored affinity). Right: DiffDock confidence vs. GNINA minimized affinity (*r* = 0.036), showing weak correlation, reflecting conformational changes introduced during post-docking energy minimization.

**Fig. 6.**
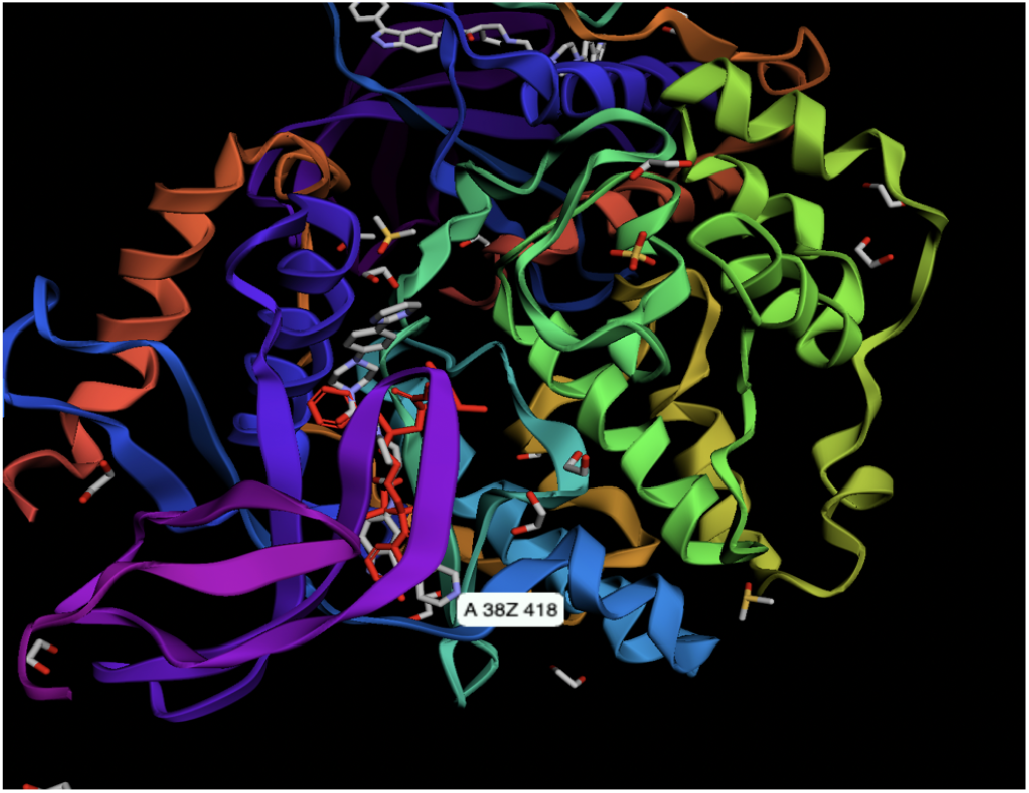
3D protein-ligand docking interaction view for the top-ranked Ribavirin pose against the Zika NS5 polymerase (PDB: 4QTB; DiffDock confidence −1.4; GNINA minimized affinity −7.31). Binding interactions at the NS5 active site include hydrogen bonds, hydrophobic contacts, and *π*–*π* stacking, highlighting residues critical to viral replication.

### Multi-metric ligand similarity scoring supports drug candidate triage

Following docking, AGPI’s ligand similarity module computes pharmacological similarity between candidate compounds using multiple molecular fingerprint metrics. For the Zika case study, two candidate ligands were profiled (SMILES strings provided in Supplementary Note 3). Similarity scores are reported in Table 2 and visualized in Figure 7.

**Table 2.** Ligand similarity scores for two candidate Zika therapeutic compounds across four molecular fingerprint metrics.

| Metric | Score | Interpretation |
| --- | --- | --- |
| Tanimoto (Morgan) | 0.1951 | Low structural similarity |
| Dice (Morgan) | 0.3265 | Moderate feature overlap |
| Cosine (Morgan) | 0.3266 | Moderate feature overlap |
| Tanimoto (MACCS) | 0.3599 | Moderate structural similarity |

**Fig. 7.**
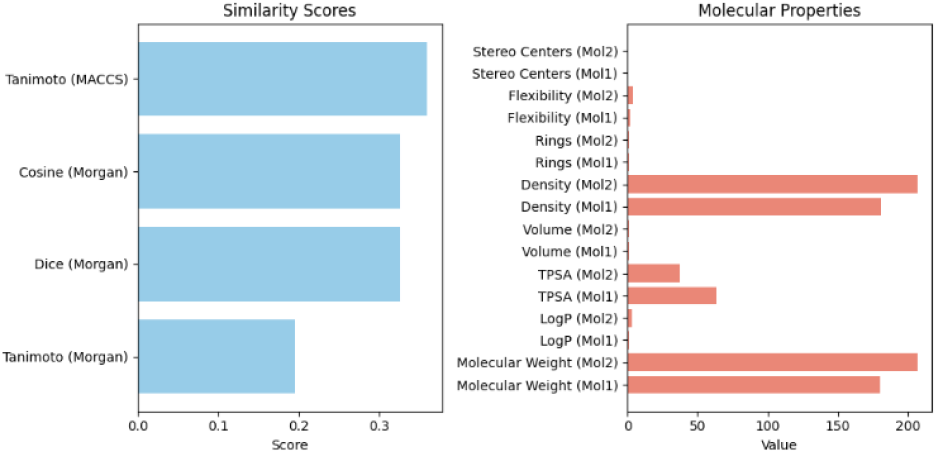
Ligand similarity and molecular property comparison for two candidate Zika therapeutic compounds. Left: similarity scores computed using Tanimoto (MACCS, Morgan), Cosine (Morgan), and Dice (Morgan) fingerprints. Right: molecular property profiles including stereo centres, ring counts, TPSA, LogP, and molecular weight. Distinct profiles across all metrics indicate different pharmacological behaviours and likely ADMET properties.

The two compounds exhibit distinct pharmacological profiles across all four metrics. Differences in polarity and hydrophobicity suggest divergent ADMET (absorption, distribution, metabolism, excretion, and toxicity) profiles, which would be expected to produce different binding behaviours and therapeutic windows. Multi-metric similarity scoring of this kind provides a principled basis for triaging drug candidates prior to resource-intensive experimental validation, and could guide selection of structural analogues for lead optimization.

## Discussion

AGPI demonstrates that a unified, AI-guided genomic pipeline can deliver accurate, interpretable, and clinically relevant insights spanning the full spectrum of pathogen analysis—from sequence-level classification to in silico therapeutic prioritization. The hybrid Conv1D–BiGRU architecture proved well-suited to genomic sequence modeling: convolutional layers efficiently extract local sequence motifs, while the bidirectional GRU captures long-range dependencies that are biologically important but invisible to uni-directional or purely local models. The near-perfect validation accuracy (99.61%) warrants careful interpretation: it likely reflects both the genuine discriminability of the 40 pathogen classes at the DNA sequence level and the relatively controlled nature of the NCBI-retrieved dataset, where sequences are well-annotated and taxonomically curated. The systematic improvement from 80.50% to 94.90% through iterative held-out evaluation confirms the value of structured, misclassification-targeted fine-tuning as a practical strategy for improving model robustness on diverse sequence inputs.

A direct quantitative comparison between AGPI and existing genomic classifiers such as DeepVirFinder (2), VirNet (3), and transformer-based genomic models including DNABERT (7) was not undertaken because these systems differ substantially in scope, training data, taxonomic coverage, and evaluation protocols. DeepVirFinder and VirNet were developed primarily for viral sequence identification from metagenomic datasets, whereas AGPI was designed as a multi-class platform spanning viruses, bacteria, fungi, and protozoan pathogens while integrating downstream biological enrichment, structural visualization, and therapeutic target prioritization. Consequently, numerical performance comparisons across independently curated datasets would not provide a fair or directly interpretable benchmark. Instead, AGPI should be viewed as a complementary end-to-end framework that unifies multiple stages of pathogen analysis within a single interpretable workflow. Future studies employing standardized benchmark datasets and identical train-test splits across competing methods will enable rigorous quantitative comparisons under equivalent experimental conditions.

The integration of DiffDock (4) for molecular docking represents a meaningful methodological advance over traditional rigid-body approaches such as AutoDock Vina (17). Diffusion-based docking explicitly models protein-ligand conformational flexibility, yielding more physically realistic binding poses. Importantly, correlation analysis across 20 docking poses revealed a moderate relationship between DiffDock confidence and GNINA scored affinity (*r* = − 0.522) but a weak relationship with GNINA minimized affinity (*r* = 0.036), indicating that these two scoring functions are complementary rather than redundant.

The 3D binding analysis of the top-ranked Ribavirin pose— revealing interactions at the NS5 polymerase active site via hydrogen bonds, hydrophobic contacts, and *π*–*π* stacking— should be interpreted as an in silico hypothesis requiring experimental validation rather than evidence of therapeutic efficacy. Ribavirin is a guanosine analogue with well-documented broad-spectrum antiviral activity that has been investigated against numerous RNA viruses through mechanisms including inhibition of viral RNA synthesis and modulation of nucleotide metabolism. Because the NS5 RNA-dependent RNA polymerase is essential for Zika virus genome replication, its identification as the predicted binding target provides a biologically plausible rationale for prioritizing Ribavirin within AGPI’s therapeutic screening pipeline. Although the docking scores and interaction profiles suggest favorable binding, these computational predictions should be regarded as hypothesis-generating and require validation through biochemical assays, cell-based infection models, and ultimately in vivo studies before any conclusions regarding therapeutic efficacy can be drawn.

The multi-metric ligand similarity framework addresses a recognized weakness of single-fingerprint screening: no single molecular fingerprint captures all pharmacologically relevant structural features (6). By combining Tanimoto, Dice, and Cosine similarities across both Morgan and MACCS fingerprints, AGPI provides a richer basis for compound comparison. The divergent scores observed between the two Zika candidate ligands—with Tanimoto (Morgan) of only 0.1951 but Tanimoto (MACCS) of 0.3599—illustrate how different fingerprint methods weight structural features differently, and how relying on a single metric could either over-or underestimate pharmacological similarity. Multi-metric triage before committing to in vitro validation is therefore both scientifically more rigorous and more cost-effective.

Several limitations warrant acknowledgment. **Sequence-identity-based data partitioning** represents the most important methodological limitation of the current evaluation. Although exact-match deduplication was applied prior to splitting, sequences from the same species or closely related strains may share substantial nucleotide similarity, potentially inflating generalization estimates. The near-perfect validation accuracy (99.82%) should therefore be interpreted with caution: it likely reflects in part the absence of identity-controlled splitting rather than solely the model’s ability to generalize to genuinely novel sequences. Future work should apply CD-HIT (10) or equivalent clustering tools to enforce a maximum pairwise sequence identity threshold (e.g., 90%) across training, validation, and held-out evaluation splits. This would yield a more conservative and clinically meaningful estimate of generalization performance, particularly for deployment scenarios involving emerging variants or closely related pathogen strains not represented in the training corpus.

Fixed-length sequence input (1,000 nucleotides) may reduce performance on very long or highly fragmented genomic sequences. The training corpus of 40 pathogen classes does not represent the full breadth of clinically encountered pathogens, and expanded datasets including emerging variants will be required for broader surveillance deployment. All docking predictions are in silico and require experimental validation—GNINA minimized affinity values are informative for ranking but are not directly comparable to measured binding constants (*K*_*d*_, IC_50_). Finally, the biological enrichment corpus is static; integration with live literature APIs such as PubMed Entrez or Europe PMC would keep recommendations current with emerging research.

Beyond its predictive performance, AGPI illustrates the value of integrating complementary AI methodologies into a unified decision-support framework for genomic analysis. Rather than treating pathogen classification, structural modeling, literature-based biological enrichment, and therapeutic prioritization as isolated computational tasks, the platform links these analyses into a single interpretable workflow that mirrors the progression of biological investigation from sequence identification to hypothesis generation. Such integration has the potential to accelerate exploratory analyses during infectious disease outbreaks by enabling researchers to rapidly contextualize newly sequenced pathogens and identify candidate therapeutic targets for downstream experimental evaluation. Although AGPI is not intended to replace laboratory validation or clinical decision-making, it demonstrates how explainable AI-driven pipelines can complement existing bioinformatics resources by reducing analytical fragmentation and facilitating hypothesis-driven research. These capabilities align with broader trends in high-performance medicine, where artificial intelligence is increasingly used to augment clinical and translational decision-making by integrating multimodal biological data into actionable insights (18).

Future development priorities include: real-time genome scanning via streaming FASTA input; expanded training datasets covering emerging pathogens; sequence identity-controlled data splitting; integration of AlphaFold2 (19) and Rosetta for complementary docking strategies; incorporation of DNABERT (7) for enhanced genomic sequence representation; automated lead optimization workflows; and cloud-based collaborative research dashboards. Connecting AGPI to healthcare APIs and global surveillance networks such as GISAID could create a dynamic knowledge loop, substantially improving real-time clinical utility for emerging infectious disease response.

## Conclusions

We present AGPI, a unified AI-powered platform that addresses a critical gap in genomic bioinformatics by integrating pathogen classification, biological enrichment, interactive 3D visualization, and protein-ligand docking into a single, interpretable pipeline. The platform’s hybrid Conv1D– BiGRU classifier achieves 99.61% validation accuracy across 40 pathogen classes and 94.90% accuracy on 600 randomized sequences after iterative refinement. End-to-end demonstration on Zika virus confirmed both the diagnostic accuracy of the classification module (96.14% confidence) and the therapeutic utility of the docking module, with Ribavirin identified as a top repurposing candidate against the NS5 polymerase. Multi-metric ligand similarity scoring provided additional pharmacological discrimination between candidate compounds. By making complex genomic workflows accessible without sacrificing scientific rigour, AGPI contributes to the broader goal of AI-augmented clinical genomics and represents a scalable foundation for real-time pathogen surveillance and drug discovery.

## Data Availability

The code and data supporting this study are available from the corresponding author upon reasonable request.

## ACKNOWLEDGEMENTS

The authors thank CMU’s College of Engineering and VIT for institutional support. The open-source communities behind Biopython, DiffDock, GNINA, and NVIDIA EsmFold are gratefully acknowledged.

## Methods

### Data collection

DNA sequences for 40 pathogen classes were retrieved from the NCBI database using BLASTn with default parameters, exported in FASTA format. Selection criteria were: (i) clinical relevance to global disease burden; (ii) diversity of geographic and biological origin; and (iii) representation across major viral, bacterial, and fungal families, including Zika, Dengue, Ebola, SARS-CoV-2, Hepatitis, Malaria, and *Vibrio cholerae*, among others.

### Preprocessing

Sequences were loaded from preprocessed CSV files. Duplicates were removed and unique integer class labels assigned per pathogen. Nucleotide bases were one-hot encoded (A, T, C, G → 4-dimensional binary vectors). All sequences were truncated or zero-padded to 1,000 nucleotides to ensure uniform input dimensionality.

### Model architecture and training

The classifier employs a hybrid Conv1D–BiGRU architecture designed to capture both local nucleotide motifs and long-range sequence dependencies. DNA sequences were first converted into one-hot encoded representations. For a nucleotide sequence of length *L*, each nucleotide was represented as a four-dimensional binary vector,

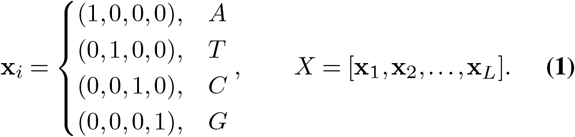

The encoded sequence was passed through one-dimensional convolutional filters to identify local sequence motifs,

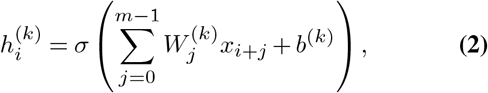

where *m* denotes the kernel size, *W* ^(*k*)^ the *k*th convolutional filter, *b*^(*k*)^ the corresponding bias, and *σ*(*·*) the ReLU activation function. Max-pooling and dropout were subsequently applied for dimensionality reduction and regularization.

The resulting feature maps were processed by a bidirectional gated recurrent unit (BiGRU), which models contextual dependencies in both forward and reverse sequence directions.

The hidden state update is defined as

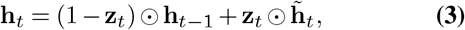

where **z**_*t*_ denotes the update gate, 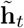 the candidate hidden state, and ⊙ element-wise multiplication. The forward and backward hidden states were concatenated before classification.

The final dense layer produced class probabilities through the Softmax function,

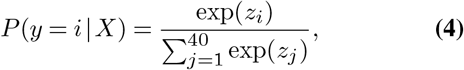

where *z*_*i*_ represents the output logit for pathogen class *i*. Model parameters were optimized using categorical cross-entropy loss,

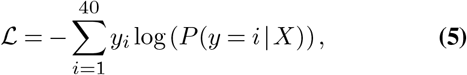

where *y*_*i*_ denotes the one-hot encoded ground-truth label. Optimization was performed using the Adam optimizer with early stopping based on validation accuracy to reduce overfitting. The principal architectural and optimization hyperparameters are summarized in Table 3.

**Table 3.** Hyperparameters used for training the AGPI Conv1D–BiGRU classifier.

| Parameter | Value |
| --- | --- |
| Input sequence length | 1,000 nucleotides |
| Input representation | One-hot encoding (4 channels) |
| Conv1D Layer 1 | 27 filters, kernel size = 4 |
| Activation (Conv1D) | ReLU |
| MaxPooling1D | Pool size = 3 |
| Dropout | 0.20 |
| Conv1D Layer 2 | 14 filters, kernel size = 2 |
| BiGRU units | 128 |
| Dense layers | 128, 64, 64, 16 neurons |
| Dense activation | ReLU |
| Output layer | 40 neurons (Softmax) |
| L2 regularization | $\lambda = 0.01$ (final hidden layer) |
| Loss function | Categorical cross-entropy |
| Optimizer | Adam |
| Batch size | 128 |
| Maximum epochs | 500 |
| Early stopping monitor | Validation accuracy |
| Minimum improvement ($\Delta$) | 0.0005 |
| Patience | 8 epochs |
| Restore best weights | Yes |
| Evaluation metric | Accuracy |

**Table 4.** SMILES strings for candidate ligands used in the Zika therapeutic triage analysis.

| Compound | SMILES |
| --- | --- |
| Molecule 1 | CC(=O)OC1=CC=CC=C1C(=O)O |
| Molecule 2 | CC(C)CC1=CC=C(C=C1)C(C)C(=O)O |

The BiGRU layer employed a ReLU activation function as implemented in the training architecture. All remaining GRU gating operations retained the standard Keras implementation.

### Iterative held-out evaluation

Classification accuracy was assessed on 600 held-out sequences (15 per class, 40 classes), drawn after train/validation splitting was finalized. Each sequence was evaluated independently. Four rounds of evaluation were conducted with iterative fine-tuning applied to misclassified samples between rounds. Final accuracy was reported at convergence (Round 4).

### Biological enrichment

Following classification, the enrichment module retrieves clinical symptoms, virulence factors, immune evasion proteins, and recommended PDB structures and PubChem compounds from a curated corpus of 245 peer-reviewed studies. The corpus was assembled manually by the author, with studies selected from PubMed on the basis of direct relevance to the 40 pathogen classes included in the training dataset; one or more representative studies were identified per pathogen, prioritizing primary research articles describing clinical presentation, virulence mechanisms, and known therapeutic targets.

### 3D genome visualization

Protein structures are predicted from user-input DNA sequences using NVIDIA EsmFold (14). Annotation and interactive rendering are implemented with Biopython (15). The genome browser supports clickable annotation of coding and functional regions at the residue level.

### Protein-ligand docking

DiffDock (4) performs diffusion-based protein-ligand docking. Users specify a PDB ID for the target protein and either a SMILES string or PubChem CID for the ligand; PubChem CIDs are automatically resolved to SMILES via the PubChem REST API. ESM embeddings are used to pre-process protein sequences for DiffDock input. Twenty binding poses are generated per run, scored by Diff-Dock’s internal confidence metric and independently evaluated using GNINA for raw and minimized binding affinities. Results are presented in ranked tabular format and available for download as a compressed archive.

### Ligand similarity scoring

Post-docking, candidate ligands are compared using four molecular fingerprint metrics: Tanimoto (Morgan and MACCS fingerprints), Dice (Morgan), and Cosine (Morgan). Molecular property profiles—stereo centres, ring counts, density, volume, TPSA, LogP, and molecular weight—are additionally computed and visualized to contextualize pharmacological differences between compounds.

---

**Supplementary Note 1: Extended dataset statistics**

The 40 pathogen classes used for training and evaluation span three clades: viruses (including RNA viruses such as Zika, Dengue, Ebola, SARS-CoV-2, and Hepatitis, and DNA viruses), bacteria (gram-positive and gram-negative, including *Vibrio cholerae*), protozoan parasites (including *Plasmodium* spp., responsible for malaria), and fungi. Classes were selected to ensure both clinical prevalence and taxonomic diversity, avoiding over-representation of any single viral or bacterial family. All sequences were retrieved in FASTA format from NCBI GenBank using BLASTn queries with default parameters and reviewed for annotation quality prior to inclusion in the training corpus.

**Supplementary Note 2: Docking simulation parameters**

For the Zika NS5 polymerase docking simulation, the target protein was specified by PDB ID 4QTB. Ribavirin was retrieved by PubChem CID, with the corresponding SMILES string automatically resolved via the PubChem REST API. ESM embeddings were computed for the protein sequence prior to DiffDock inference. Twenty binding poses were generated; each was scored by DiffDock’s confidence metric and independently re-scored using GNINA with minimized affinity. The top-ranked pose (DiffDock confidence − 1.4; GNINA minimized affinity − 7.31) was selected for structural visualization and interaction analysis.

**Supplementary Note 3: Candidate ligand SMILES strings**

SMILES strings for the two candidate Zika therapeutic compounds profiled in the ligand similarity analysis are provided below.

